# Modulation of large-scale cortical coupling by transcranial alternating current stimulation

**DOI:** 10.1101/484014

**Authors:** Bettina C. Schwab, Jonas Misselhorn, Andreas K. Engel

## Abstract

**Background:** Long-range functional connectivity in the brain is considered fundamental for cognition and is known to be altered in many neuropsychiatric disorders. To modify such coupling independent of sensory input, noninvasive brain stimulation could be of utmost value.

**Objective:** First, we tested if transcranial alternating current stimulation (tACS) is able to influence functional connectivity in the human brain. Second, we investigated the specificity of effects in frequency and space.

**Methods:** Participants were stimulated bifocally with high-definition tACS in counterbalanced order (1) in-phase, with identical electric fields in both hemispheres, (2) anti-phase, with phase-reversed electric fields in the two hemispheres, and (3) jittered-phase, generated by subtle frequency shifts continuously changing the phase relation between the two fields. EEG aftereffects were analyzed systematically in sensor and source space.

**Results:** While total power and spatial distribution of the fields were comparable between conditions, global pre-post stimulation changes in EEG connectivity were larger after in-phase stimulation than after anti-phase or jittered-phase stimulation. Those differences in connectivity were restricted to the stimulated frequency band and decayed within the first 120 s after stimulation offset. Source reconstruction localized the maximum effect between the stimulated occipitoparietal area.

**Conclusion:** The relative phase of bifocal alpha-tACS modulated alpha-band connectivity between the targeted regions. As side effects are not expected to differ between the stimulation conditions, we conclude that neural activity was phase-specifically influenced by the electric fields. We thus suggest bifocal high-definition tACS as a tool to manipulate long-range cortico-cortical coupling which outlasts the stimulation period.

## 1. Introduction

Oscillatory activity and its synchronization at various temporal and spatial scales are regarded as crucial for cognition and behavior. In particular, oscillatory coupling has been suggested to functionally link remote brain areas by synchronously opening windows for communication [1, 2]. During sensory stimulation, large-scale connectivity, as measured by EEG or MEG, may thus form task-related networks within cortex [1, 3, 4]. But also ongoing resting-state connectivity, emerging from brain-intrinsic factors rather than external stimuli, was hypothesized to reflect physiological brain function and to bias processing of upcoming stimuli [2].

Due to their prominence in the resting-state EEG, *α*-oscillations and connectivity in the *α*-band have been studied extensively. *α*-oscillations were typically related to functional inhibition [5] and were proposed to route information to task-relevant regions [6]. Accordingly, phase synchronization of *α*-oscillations may have a crucial role in orchestrating the exchange of information in the brain [7] relevant for attention, memory, and executive functioning [5, 8]. Reduced or increased *α*-coupling in disease could therefore be related to cognitive impairment [9, 10, 11].

Despite the many studies demonstrating correlations between *α*-coupling and aspects of cognition, it is hard to find evidence for a causal relation based on EEG or MEG measurements alone. In contrast, noninvasive brain stimulation might be able to directly interfere with cortical dynamics. Especially transcranial alternating current stimulation (tACS), applied at different brain sites, was suggested to frequency-specifically modulate coupling between cortical areas [15, 16, 17]. tACS could thus be a valuable tool to investigate the role of functional connectivity for cognition, and, in later stages, to re-adapt pathological coupling in patients towards a physiological level.

Yet, up to now it is unclear what the immediate as well as outlasting effects of tACS on cortical electrophysiology are, and, critically, whether they are specific to the stimulated frequency band and cortical area. While *α*-tACS can lead to an increase in *α*-power after the offset of stimulation [18, 19, 20, 21], much less is known about modulation of coupling. Behavioral changes under tACS have been described and were in some studies related to connectivity modulation [15, 16, 17, 21, 22, 23]. However, so far, conclusive evidence for connectivity modulation is missing, as the analysis of EEG and MEG data is impeded by a large, hard-to-predict stimulation artifact [24, 25]. Connectivity analyses of stimulation-outlasting effects were often confounded by power effects or were restricted to sensor level analyses limiting spatial resolution.

Hence, we present an in-depth investigation of cortical large-scale synchronization after bifocal *α*-tACS. To bypass the stimulation artifact, we analyzed resting-state EEG data before and after tACS (Fig. 1A), with conditions differing in the phase relation of the applied fields (Fig. 1B). In particular, all tACS conditions involved *α*-stimulation with comparable power distributions of the electric fields (Fig. 1C), avoiding different aftereffects in *α*-power, different stimulation of peripheral nerves, and different retinal stimulation. We were able to frequency-specifically modulate functional *α*-coupling between the stimulated regions in a time window 0-100 s after tACS offset, opening up the possibility to use tACS as a powerful manipulator of functional connectivity.

## 2. Methods and Materials

### 2.1. Experimental setup

#### 2.1.1. Participants

28 participants entered the study and received financial compensation. One participant was excluded as she closed her eyes during parts of the recording, and technical problems led to the exclusion of another participant. Since two other participants reported discomfort during test stimulation, no recordings were obtained from them. Hence, we were able to complete data collection from 24 participants in a counterbalanced sequence of three stimulation conditions. These final participants (12 male, 12 female) were on average 26 ± 4 years old. Vision was normal or corrected to normal and none of them had a history of neurological or psychiatric disorders. Participants gave written informed consent after introduction to the experiment. The ethics committee of the Medical Association Hamburg approved the study.

**Figure 1:**
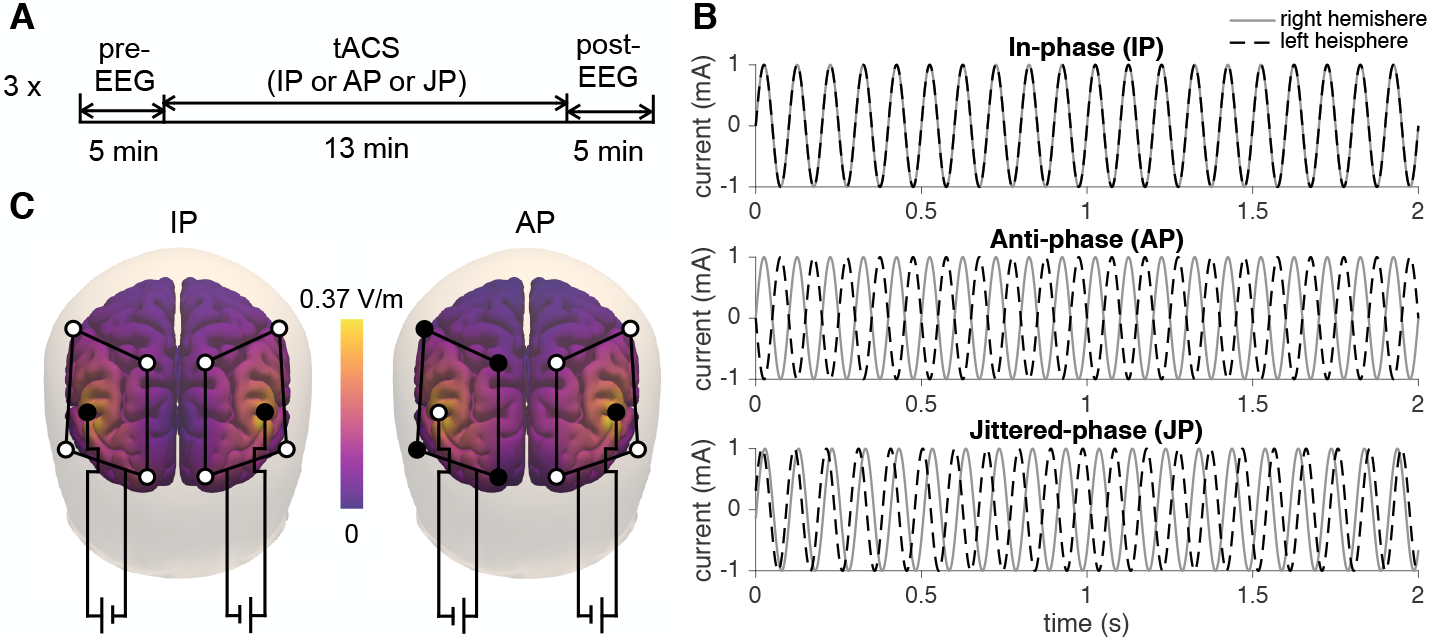
Experimental setup. A: Timeline of resting-state recordings. EEG and ECG were recorded in the eyes open state before and after tACS in three sessions. Pupil dilation was recorded during the whole time course. The sequence of tACS conditions was counterbalanced between participants. B: Stimulation current waveforms of the three tACS conditions, applied on the right hemisphere (gray) and left hemisphere (black dashed), shown for 2 s. C: Peak power, for example at time point 25ms, of bifocal occipito-parietal stimulation fields for IP and AP stimulation. Magnitudes of the applied electric fields were similar for IP and AP stimulation. In contrast, the direction of the fields differed between conditions, with IP stimulation generating identical fields within each hemisphere and AP stimulation generating fields of opposite direction. See Supplementary Material A for details.

#### 2.1.2. Electrophysiological recordings

Resting-state recordings included electroencephalogram (EEG) and electrocardiogram (ECG). Participants were seated comfortably in a dimly lit, electrically shielded room and asked to fixate a cross displayed on the screen. 64 Ag/AgCl EEG electrodes mounted on an elastic cap (Easycap) were prepared with an abrasive conducting gel (Abralyt 2000, Easycap) to keep impedances below 20 *k*Ω. EEG signals were referenced to the nose tip and recorded using BrainAmp amplifiers (Brain Products GmbH). ECG was measured as the voltage between right infraclavicular fossa and left flank. All participants took part in three sessions on one day. Each session was 23 min long: After a period of 5 min EEG resting-state recording, tACS was applied for 13 minutes, followed by 5 min post-tACS EEG recording (Fig. 1A). Between the sessions, a break of at least 5 min was held.

#### 2.1.3. Transcranial stimulation

Ten additional Ag/AgCl electrodes with 12 mm diameter were mounted between EEG electrodes for application of tACS. After preparation with Signa electrolyte gel (Parker Laboratories Inc), impedances of each outer electrode to the center electrode were allowed to range between 1 and 120 kΩ. Subsequently, the four outer electrodes of each montage were electrically connected to each other and to an alternating current source (DC-Stimulator plus, NeuroConn), with the center electrode as the reference (see Fig. 1C). For each montage, similar impedances were attempted (differences to average impedance below 20%), leading to a total impedance per montage below 20 k*Ω*. Participants were made familiar with tACS by slowly increasing the amplitude of a test stimulus. During each session, *α*-tACS was applied with a peak-to-peak amplitude of 2 mA for 13 min, including a linear ramp-up within the first 10 s. The signal for stimulation was computed in MATLAB and produced by a NI-DAQ device. The three *α*-tACS conditions were in-phase (IP) stimulation, anti-phase (AP) stimulation, and jittered-phase (JP) stimulation. Participants were blinded towards those conditions. IP and AP stimulation consisted of two 10 Hz sinusoidal currents with phase offset zero and *π*, respectively, applied at the two hemispheres. For JP stimulation, the two stimulating currents independently changed their frequency between 9.5 and 10.5 Hz at constant amplitude (Fig. 1B). While spectral power of all stimulating currents was focused around 10 Hz, coherence between the two signals was low in the JP condition (see Supplementary Material B). Hence, all three conditions involved *α*-stimulation, but differed in the phase relation between the stimulating currents. The sequence of tACS conditions was counterbalanced between participants.

#### 2.1.4. tACS electrode montage and simulation of the applied electric field

Spatial application of tACS was chosen to target generators of *α*-activity in parieto-occipital regions. We used two 4-in-1 montages similar to Helfrich et al. [17]. At each hemisphere, one montage was applied and connected to a stimulator, as shown in Fig. 1C. The use of separate stimulators for each montage ensured a focal field underneath the stimulation electrodes.

In order to quantify and compare field distributions for the different stimulation conditions, we simulated the electric field induced by tACS. The leadfield matrix *L* was computed with the boundary element method volume conduction model by Oostenveld et al. [26] and segmented using the AAL atlas, equally to source reconstruction of EEG signals. Distances between grid points were 1 cm in all three spatial dimensions. The electric field 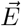 at location 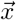 could then be calculated by linear superposition of the evoked fields of all injected currents *α*_*i*_ at stimulation electrodes *i* = {*1*,*2*, …,*10*} as

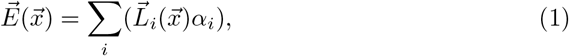

with the field strength 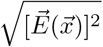. Contrasts in field strength were either computed as vectorial differences (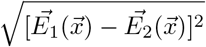 capturing differences in field direction as well as absolute strength) or absolute differences (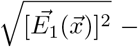 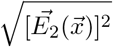; capturing differences in absolute field strength only).

To additionally simulate the stimulating field with a higher spatial precision, the leadfield matrix was also computed with the boundary element method on a volume conduction three-shell model which was reconstructed from the MNI template brain (see Fig. 1C). Here, spatial resolution was 5 mm in all three dimensions and allowed for detailed description of the stimulating fields (Supplementary Material A).

### 2.2. Eye tracking and questionnaire on peripheral sensations

During the whole time course of recording sessions, pupil dilation and its confidence (∈ [0, 1], with value one indicating highest confidence) were recorded for both eyes with head-mounted eye trackers (Pupil Labs, Germany) at 60 Hz. For one participant, technical problems did not allow for recording of pupil dilation in the third session (JP stimulation). After the final session, participants were asked for their subjective sensation of tACS and differences between sessions in a qualitative questionnaire including the following questions:

- When during the session did you feel any electric stimulation?
- Please describe the sensation on your skin during stimulation.
- Did you have other sensations during stimulation, for example visual sensations?
- Did you feel any differences between the three sessions?

### 2.3. Data processing

#### 2.3.1. EEG pre-processing

64-channel EEG data, recorded at 5 kHz, was pre-processed in MATLAB 2015b (The MathWorks Inc) and FieldTrip [27]. Time series were segmented into 300 s trials before (“pre recording”) and after tACS (“post recording”) prior to preprocessing to avoid leakage of the tACS artifact into the pre and post recordings. The first 500 ms of each trial were discarded as post recordings may contain remaining tACS artifacts related to capacitive effects within this period. Trials were re-referenced to common average, two-pass high-pass (1 Hz) as well as low-pass (25 Hz) filtered with a second order Butterworth filter, and down-sampled to 100 Hz.

For the six combined trials of each participant, an independent component analysis (ICA) using the infomax ICA algorithm [28] was computed over all channels. Pearson’s linear correlation coefficient [29] between the largest 30 independent components (ICs) and the ECG as well as the two electrooculogram (EOG) traces were computed. ICs that showed a correlation coefficient larger than 0.2 to the ECG or one of the EOGs were removed in order to minimize cardiac and eye-blink artifacts. After back-projection to sensor space, channels were screened for high noise levels. If the standard deviation of a channel was higher than three times the median standard deviation of all channels, the channel was excluded. On average, 4.7 ± 1.5 components (mean ± standard deviation) and 1.1 ± 1.5 channels per participant were removed. Both exclusion of ICs and channels did thus not differ between conditions as exactly the same ICs and channels were removed. The whole pre-processing pipeline was automated and did not rely on subjective decisions of the investigator.

#### 2.3.2. Source reconstruction

Exact low resolution brain electromagnetic tomography (eLORETA) [30, 31] - a discrete, three-dimensional distributed, linear, weighted minimum norm inverse solution - was used to estimate neural activity at source level. Before source projection, we band-pass filtered sensor data with a 2nd order butterworth filter between 7 and 13 Hz. eLORETA was then applied with 1% regularization, using the boundary element method volume conduction model by Oostenveld et al. [26]. Three-dimensional time series of dipoles were reconstructed at a linearly spaced grid of 1074 cortical and hippocampal voxels with distance 1 cm in all three spatial directions, and regions were assigned following the Automated Anatomical Labeling (AAL) atlas parcellation [32]. Spatiospectral decomposition (SSD) [33], maximizing activity between 9-11 Hz while suppressing activity in the flanking bands 8-9 Hz and 11-12 Hz, was applied to time series of the three spatial dimensions at each voxel. To reduce the dimensionality of source space data, only the largest component was considered for further analysis [34]. As this approach leads to a specific filtering of activity around 10 Hz, source space effects were only studied within the frequency band 9-11 Hz.

#### 2.3.3. Time-frequency analysis

EEG data were analyzed in sensor as well as in source space. At each electrode in sensor space and each voxel in source space, power in frequency bands of 2 Hz width was computed within 1 s segments without overlap and subsequently averaged over the 100 s window of interest. Likewise, for all electrode pairs in sensor space and all voxel pairs in source space, the absolute value of the imaginary part of coherency [35], also termed imaginary coherence, was computed as

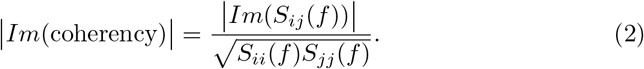

The cross- and auto-spectra 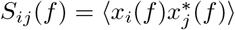 were estimated from Fourier transforms of 1 s EEG time series of channel i and j, *x*_*i*_(*f*) and *x*_*j*_(*f*), respectively, and averaged over the 100 s window of interest. To keep the time distance between pre- and post-recordings constant for comparisons involving a time course, 100 s time segments were cut identically from pre- and post-recordings, leading to a constant time difference of 18 min between the onsets of each time window.

#### 2.3.4. Connectivity metrics

Global connectivity shifts were assessed via the distribution of connectivity change values between electrode or voxel pairs. We computed cumulative histograms as the fraction of pairs with a certain minimum connectivity change. In sensor space, all electrode pairs were taken into account. In source space, all interhemispheric pairs of voxels between the stimulated area in the right hemisphere (“right stimulated region”) and the stimulated area in the left hemisphere (“left stimulated region”) were included. Stimulated regions were defined as regions with average differences in vectorial field strength between IP and AP stimulation above 0.08 V/m. Averaging over many pairs further minimizes spurious local functional connectivity [35], in particular in sensor space. In source space, we additionally investigated the topology of *α*-coupling changes. Differences in imaginary coherence were computed for all pairs of voxels and averaged within regions of interest.

#### 2.3.5. Permutation statistics

To test for significance of condition differences between IP, AP, and JP stimulation, we performed permutation statistics. 100 s epochs of both pre- and post-recording were cut into trials of 1 s each. For each trial, conditions were shuffled to observe control time series, while pre-post pairings remained intact. As conditions were randomly mixed in these control time series, confidence intervals for connectivity differences between the conditions could be estimated. We computed 100 control time series (200 control time series in sensor space) and control connectivity measures. The 4950 (19900 for sensor space) difference values between the control connectivity measures served as control distributions for condition differences: The 2.5th and 97.5th percentile of these distributions defined the uncorrected lower and upper 95% confidence limits, respectively.

When three stimulation condition differences were compared, we applied Holm-correction to the confidence limits: If the contrast with the lowest p-value was significant after Bonferroni-correction, subsequent comparisons were adjusted for a lower number of comparisons. Figures show the confidence interval after the last comparison yielding significant results. For analyses on multiple time windows, the first time window (0-100 s after tACS offset) was relevant for Holm-correction. To additionally account for multiple comparisons of frequency bands, the confidence intervals were further shifted by a constant. This constant was chosen such that in the sum of all multiple comparisons, only 5% (or a lower fraction after Holm-correction for multiple stimulation conditions) of control time series yielded significant results. Hence, these corrected confidence intervals, shown in light gray, indicate a 5% probability of false significant results in the complete analysis.

#### 2.3.6. Pupil dilation

Only data from the eye with highest cumulated confidence was considered for further analysis. Data was first resampled to 20 Hz. Next, blink artifacts were identified by finding data points exceeding two standard deviations or sample points with confidence below 0.9. These samples and five sample points in the direct neighborhood of each artifactual sample were interpolated by spline interpolation. Finally, data was lowpass filtered (butterworth, cutoff: 4 Hz, order: 3) and cut to ± 20 s around tACS onset. Before averaging across participants, data was z-scored in order to account for different absolute levels of dilation that may result from varying camera angles and normalized to the 10 s preceding the onset of tACS ramp-in. Pupil responses were defined as average values of pupil dilation within the first 10 s of tACS with full amplitude (10 s after start of ramp-in).

## 3. Results

### 3.1. α-Coupling is modulated by the phase of bifocal high-definition tACS

Global *α*-power slightly increased over time in all three stimulation conditions (Supplementary Material C–A-C), impeding conclusions on the time course of connectivity (Supplementary Material C–D-F). Therefore, we only investigated condition differences of pre-post differences in both power and connectivity. Pre-tACS power and connectivity in the *α*-band did not differ significantly across conditions (Supplementary Material C–G-H). We hypothesized that the stimulation conditions, characterized by the phase relation of the two applied fields, differently affected functional connectivity.

To assess global connectivity at sensor level, we studied the overall distribution of change in imaginary coherence between pre- and post-stimulation intervals. Cumulative histograms, describing the fraction of pairs with a certain connectivity change or lower, comprising data from all participants and electrode pairs, were analyzed for the stimulated frequency band (9-11 Hz; Fig. 2). Such histograms have the advantage of demonstrating in which range changes occur and whether differences between two distributions were driven by outliers or by large parts of the population. Within the first 100 s after tACS offset, global pre-post change in *α*-connectivity was larger for IP stimulation than for AP stimulation, reflected in lower cumulative fractions for IP than for AP stimulation throughout the range of possible pre-post changes. JP stimulation led to intermediate connectivity changes for most pairs (Fig. 2A). Only for high positive connectivity changes, the fraction of pairs was lowest in the JP condition. In later time windows (Fig. 2B,C), condition differences already decayed.

Significance was tested for grand average condition differences in imaginary coherence change (Fig. 2D) and the Kolmogorov-Smirnov (K-S) distance between the distributions of imaginary coherence change (Fig. 2E). Both measures reached significance (*p* <0.05 after Holm-correction for 3 multiple stimulation comparisons) for the contrasts IP-AP and IP-JP within the first time window (0-100 s). As the effect lost significance already for the time window 20-120 s, we also computed the initial decay in steps of 2 s (Supplementary Material D). All measures lost significance before time window 20s-120s. We therefore conclude that changes in connectivity were restricted to the first 120 s after stimulation offset, and mainly focused around the first 20 s. Power differences between conditions, in contrast, were absent for those early time windows (inset in Fig. 2E). Only after 200 s, *α*-power in the JP condition was lower than in the IP and AP condition.

Finally, the observed effects were independent of power fluctuations and did not significantly interact with individual physiology such as the participants’ individual alpha frequency (IAF; Supplementary Material E). It can also be seen from all panels in Supplementary Material E that the effects were not driven by single participants but rather by smaller condition differences in many participants. Absolute z-scores of change in imaginary coherence were below 3 for all participants. In sum, we found transient global connectivity changes in the stimulated frequency band dependent on the stimulation condition.

### 3.2. Frequency-specificity of effects

To test for effects across different frequency bands, we computed the grand average change in imaginary coherence (Fig. 3A) as well as the K-S distance between the cumulative histograms of imaginary coherence change of two conditions (Fig. 3B). For both measures, the largest effects in contrasts IP-AP and IP-JP were observed around the stimulated 10 Hz - a frequency band free of power effects (inset in Fig. 3B). All in all, effects were focused on the stimulated *α*-frequency band, although variability and thus confidence intervals were largest in the lower *α*-band. Power effects were restricted to frequencies around 8 Hz.

### 3.3. Localization of effects in source space

As connectivity is difficult to localize at sensor level [34], we used an eLORETA source projection of the EEG data to study the spatial extent of condition differences within the first 100 s after stimulation offset. On sensor level, the contrast IP-AP showed clear connectivity effects and was not confounded by *α*-power effects (Fig. 2); hence, we focused on *α*-connectivity changes between these two conditions (Fig. 4).

Effects were most prominent between the stimulated occipito-parietal regions, marked by dashed lines (Fig. 4A; Supplementary Material F). Connectivity differences at interhemispheric pairs between the stimulated regions (black dashed box; “stimulated pairs”), were in 34.7% significant (uncorrected *p* <0.05), while this was the case for only 2.1% of all unstimulated pairs. Average values in coupling differences were about an order of magnitude larger for stimulated pairs (0.026±0.017) than for unstimulated pairs (0.0023±0.011; *p* <0.0001, two-sample t-test). Differences reached their maximum between left and right superior occipital gyrus. P-values for differences in connectivity change at stimulated pairs were additionally Holm-corrected for 49 region comparisons. After Holm-correction, four region pairs remained significant (corrected *p* <0.05; white dots in Fig. 4A).

**Figure 2:**
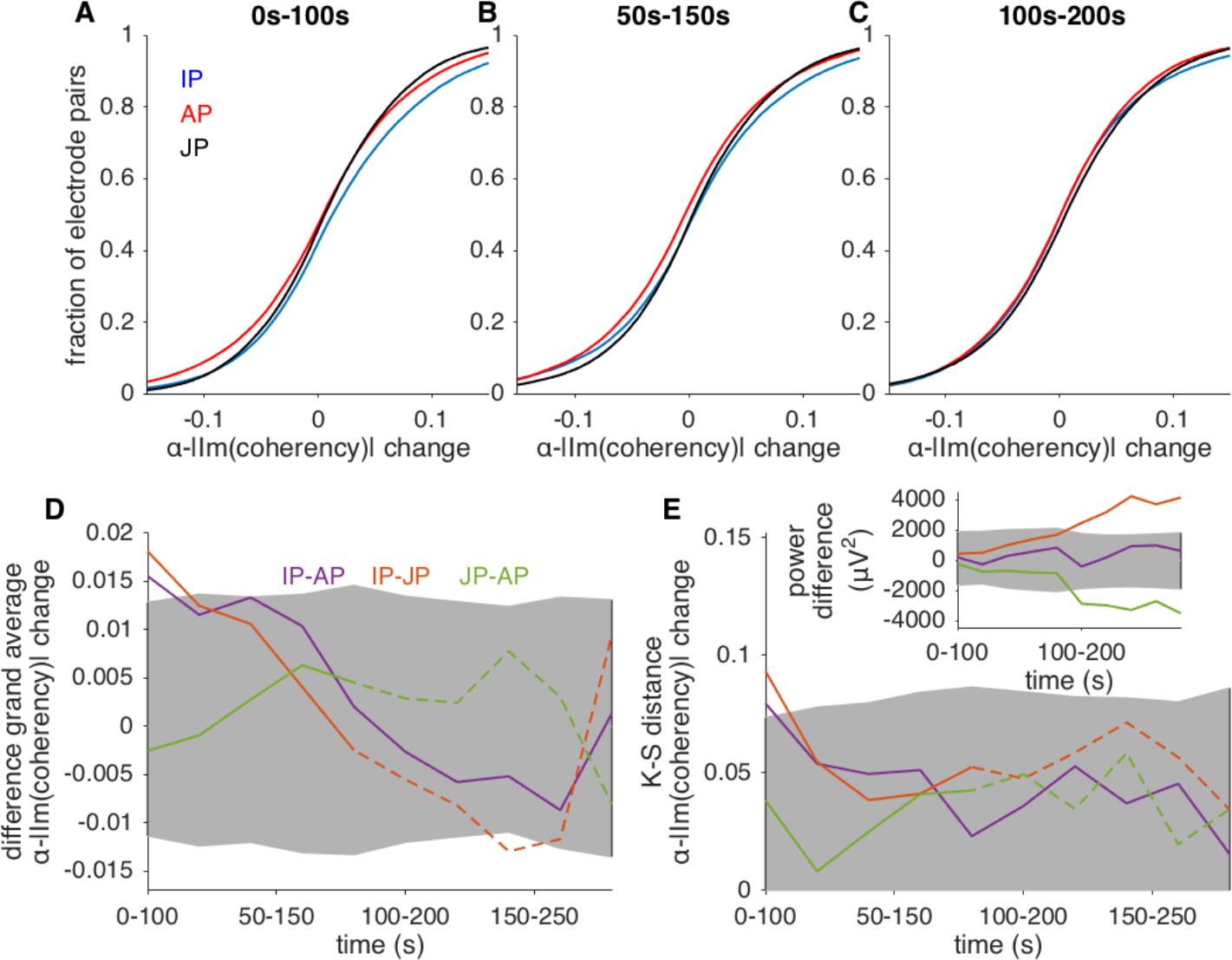
Global *α*-connectivity differs between tACS conditions. Histograms (A-C) show the cumulated fraction of *α*-imaginary coherence change from all participants and electrode pairs. Right-shifted curves thus indicate high global connectivity change compared to left-shifted curves. For the time interval 0-100s (A), IP stimulation increased connectivity compared to AP stimulation, whereas JP stimulation was associated with intermediate connectivity values. The difference decayed with time (B,C). D: Time course of grand average differences in *α*-imaginary coherence change. 95% confidence intervals Holm-corrected for multiple stimulation condition comparisons are shown in gray. The difference between IP and AP (*purple* lines) as well as between IP and JP stimulation (*orange* lines) were significant within the first time interval, 0-100 s after stimulation offset. Dashed lines indicate confounding power effects (see inset in E). E: Differences between the distributions, quantified by the K-S distance, decayed similar to grand average changes. Inset: Differences in *α*-power change between conditions. No significant differences were found for early intervals. For later intervals, JP stimulation decreased *α*-power compared to IP and AP stimulation.

**Figure 3:**
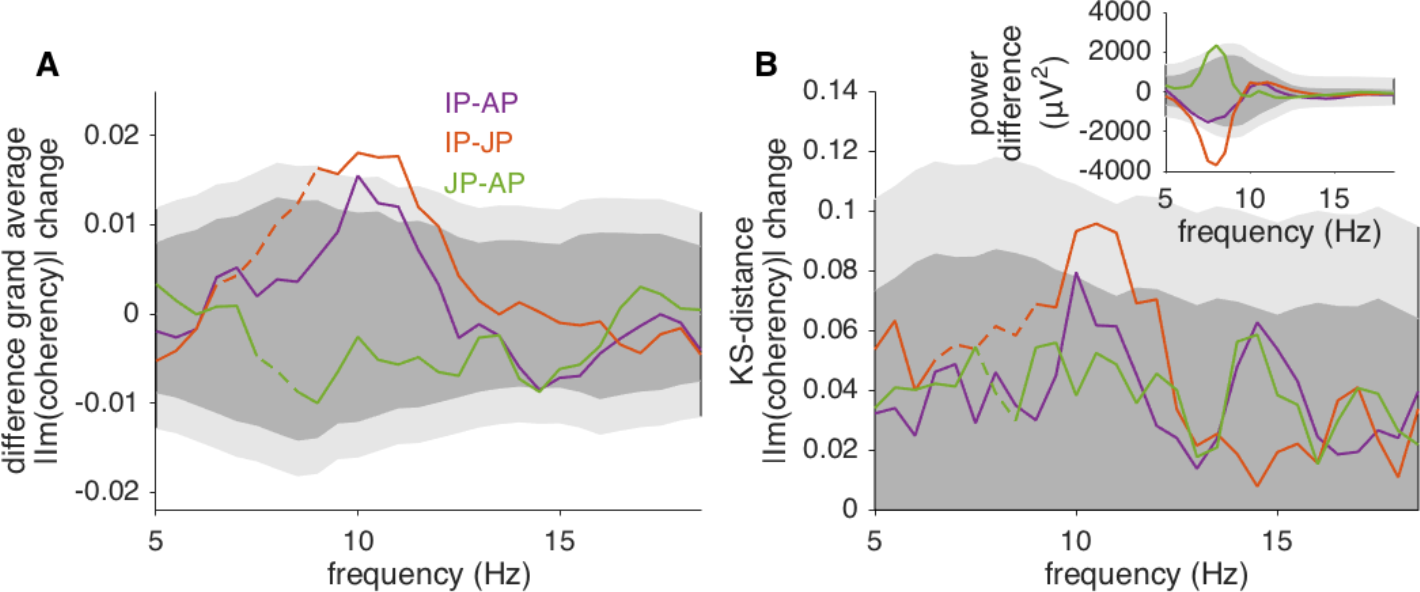
Connectivity effects in the first 100 s after stimulation offset are restricted to the stimulated frequency band around 10 Hz. A: Grand average differences in imaginary coherence for frequency bands of 2 Hz width, shifted in steps of 0.5 Hz. B: K-S distances for the same frequency bands. Dark gray: 95 % confidence intervals Holm-corrected for multiple condition comparisons. Light gray: 95 % confidence intervals additionally corrected for multiple frequency comparisons. Dashed lines indicate confounding power effects. Inset: Differences between power changes for the first 100 s after tACS offset were non-significant in the 10 Hz range.

To rule out the possibility that the observed local connectivity changes could be related to power effects, we also computed local power change between IP and AP stimulation (Fig. 4B). No significant power differences were observed, in particular also not in occipital regions, where interhemispheric coupling changes were largest. Furthermore, power effects might also influence intrahemispheric coupling, but we did not find prominent effects with the right or the left stimulated region. Thus, we do not expect power differences to drive differences in connectivity.

Looking at the population of stimulated pairs, we again computed cumulative histograms of imaginary coherence for all conditions (Fig. 4C, D). As in sensor space, IP stimulation increased connectivity compared to AP stimulation, while JP stimulation induced connectivity changes intermediate compared to the ones for IP and AP stimulation. Taken together, we found differences in connectivity change between IP and AP stimulation to be focused on the stimulated regions. Interhemispheric pairs between the stimulated regions showed similar condition differences as global sensor level data.

### 3.4. Effects spatially correlate with stimulating field

Changes in coupling between interhemispheric homologue areas were of particular interest in our study, since these areas received comparable stimulating fields in the IP condition, and comparable fields with opposite directions in the AP condition. Thus, for interhemispheric homologue regions, we further examined *α*-coupling differences (Fig. 5A), power differences (Fig. 5B), as well as vectorial and absolute differences in the stimulating fields (Fig. 5C). Note that we show average properties; peak field differences can be higher and reached 0.64 V/m for vectorial differences (Supplementary Material A).

As predicted, large changes in *α*-coupling between homologue areas were associated with large vectorial differences in field strength (Fig. 5D). In contrast, those coupling changes did not correlate with power changes (Fig. 5E), nor did the power changes correlate with vectorial differences in field strength (Fig. 5F). Therefore, we suggest phase differences in the tACS stimulus to drive changes in *α*-coupling between stimulated regions, but not *α*-power.

### 3.5. Peripheral sensations and pupil dilation related to tACS

All participants were able to detect electric stimulation in all three sessions. Typical peripheral sensations were itching, stinging, and a pulsating sensation. 18 participants reported no differences in sensation between the sessions, while the remaining six participants described the sensation to generally decrease over time with no qualitative difference between sessions. As pupil dilation at constant luminance has been linked to noradrenaline released from the locus Frontal Inf coeruleus [12] and modulates in response to sensory and noxious stimulation [13, 14], we used pupil responses as rough, but objective estimators for the strength of side effects.

**Figure 4:**
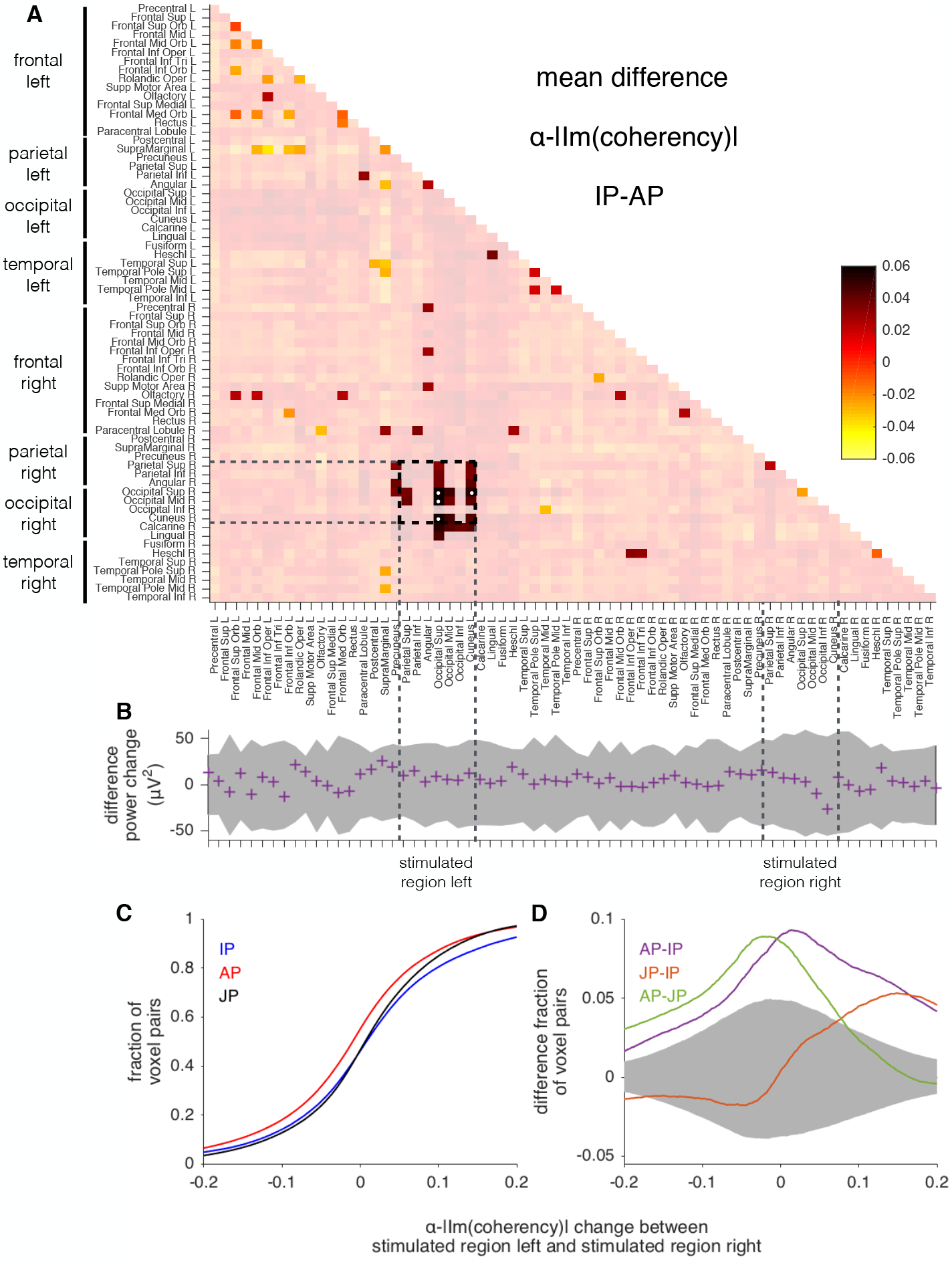
Source level characteristics of average change in *α*-connectivity within the first 100 s after stimulation offset. A: Connectivity matrix of average *α* imaginary coherence change, contrasting IP and AP stimulation conditions. Non-significant values (uncorrected *p* >0.05, permutation test) were shaded. Dashed vertical lines depict the stimulated regions, defined as an average vectorial difference in field strength above 0.08 V/m. P-values for coupling differences between the stimulated regions (black dashed box) were additionally Holm-corrected for 49 multiple regions comparisons. White dots indicate *p* <0.05 after Holm-correction. B: Differences in *α*-power change between IP and AP stimulation. No region showed significant power effects. Regions with significant coupling differences were not associated with particularly large power differences. C: Cumulative histograms of change in imaginary coherence for all pairs between the two stimulated regions. D: Difference in cumulative histograms shown in C. All condition differences extended beyond the 95% confidence limit (dark gray, Holm-corrected for three condition contrasts).

**Figure 5:**
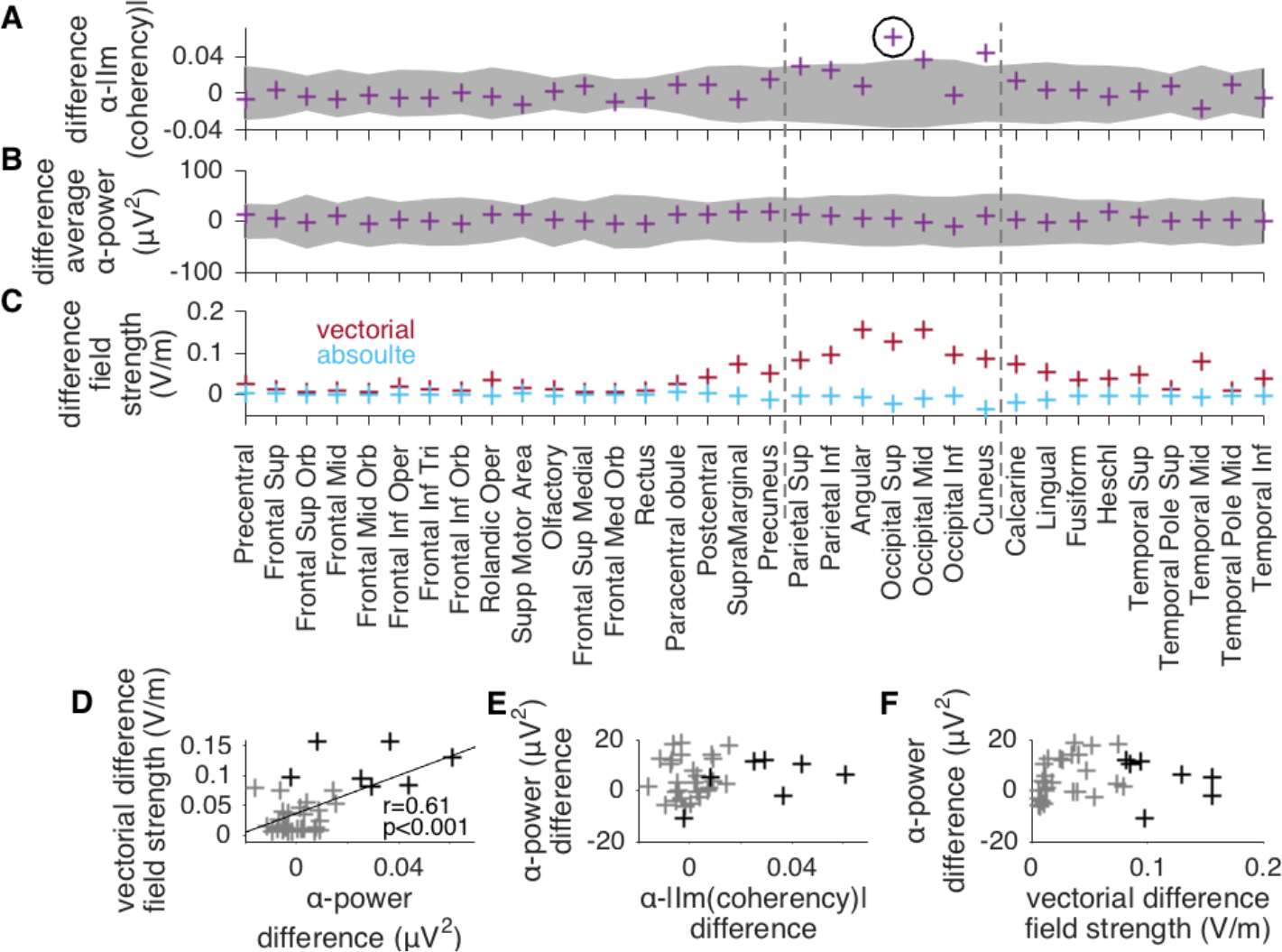
Source level change in *α*-connectivity between interhemispheric homologue areas spatially correlates with vectorial field differences. All data shown was restricted to the first 100 s after tACS offset and the contrast IP-AP. A: Changes in imaginary coherence differences between interhemispheric homologue areas for the *α*-band (9-11 Hz). Uncorrected 95% confidence intervals are shown in dark gray. Coupling between the superior occipital gyri remained significant after Holm-correction for 49 multiple region comparisons (black circle; see also Fig. 4). B: Differences in power change, averaged over interhemispheric homologue areas. C: Differences in the applied electric field between stimulation conditions IP and AP, shown as interhemispheric differences. The montage was optimized such that differences in absolute field stength (*blue* crosses) became small, while vectorial field differences were large for parietal and occipital regions (*red* crosses). The stimulated region is marked by dashed vertical lines. D: The vectorial difference in field strength correlated with change in imaginary coherence (*p* <0.001). Points representing stimulated regions are shown in black. E: In contrast, power differences did not correlate with changes in coupling. F: Power differences neither correlated with vectorial field strength, indicating the missing influence of tACS phase on *α*-power.

Pupil responses to tACS onset did not differ significantly between conditions (Supplementary Material G–A-F; uncorrected *p* > 0.5 for all condition contrasts, Wilcoxon signed rank sum test). Furthermore, there was no significant correlation between the pupil response and connectivity effects across participants (Supplementary Material G–G). Finally, five participants mentioned visual perceptions such as flickering light or phosphenes, but described them to be equally present in all three conditions. There was no significant difference in connectivity effects for those five participants compared to the remaining participants (Supplementary Material G–H; uncorrected *p* > 0.1 for all condition contrasts, Wilcoxon rank sum test).

## 4. Discussion

To our knowledge, this is the first study to describe frequency- and space-specific aftereffects of tACS on oscillatory coupling. With an optimized bifocal high-definition tACS montage and stimulation conditions differing in the phase relation of the two fields, we were able to modulate phase coupling at the stimulated frequency between the stimulated regions. For neurophysiological research, testing causal relations between coupling and behavior, our results are of high importance. Additionally, for clinical applications, the potential to enhance and prolongate effects, for example by higher stimulation amplitudes, longer duration of the stimulation, or adapted stimulation protocols, is of substantial interest.

Several preceding studies already suggested tACS as a tool to modify coupling in the brain [15, 16, 17, 21, 22, 23, 36, 37], but interpretation of connectivity in the stimulated frequency band has remained problematic. Online recordings during application of tACS [17] are subject to nonlinear stimulation artifacts which are difficult to estimate [24, 25]. While work combining tACS with fMRI circumvents those large artifacts and revealed the potential of tACS to modify coupling within the motor network, its investigation is only possible at a time scale of seconds or slower [36, 37]. Analysis of EEG aftereffects [15, 16, 17, 21, 22, 23] was so far restricted to a small number of electrodes in sensor space, where connectivity changes cannot reliably be located [34]. Importantly, none of the previous studies used conditions with stimulation differences restricted to the phases of the applied fields [38, 39]. Differences in the spatial distribution of stimulating fields may not only lead to power effects, but also to variations in tactile sensation or in the occurrence of phosphenes which may both indirectly affect neural activity. Hence, conclusions on tACS directly affecting neural activity were impeded.

Here, we optimized our study design to minimize those restrictions. We analyzed pre-post stimulation connectivity change to avoid analysis of electric artifacts during stimulation, and used detailed source reconstruction to localize effects with a centimeter resolution. Furthermore, our tACS montage yielded stimulation conditions with comparable power distributions of the electric fields, differing in their orientation. This design led to the absence of power differences between conditions at the stimulated frequency within the first 180 s after stimulation offset. The contrast IP-AP, which involved purely sinusoidal stimulation waveforms, did not induce any *α*-power differences and thus allowed for analysis of connectivity during the complete time course. Our experimental setup is therefore a major advance towards evidence for tACS modulating neural activity via electric stimulation of neural tissue.

Nevertheless, some limitations also apply to our study. First, while we demonstrate robust changes in functional connectivity, the relation of these changes to behavior was not investigated here. The physiological relevance of the observed changes in coupling is thus left to future studies combining tACS with behavioral paradigms. Second, stimulation of peripheral nerves in the skin could contribute to the obtained effects. However, a large influence of tactile sensation seems unlikely as no significant increases in coupling were observed between the postcentral gyri and other regions (see Fig. 4A). Although participants indicated no differences in sensation between the stimulation, it is a weakness of our study that we did not quantitatively rate subjective sensations. Instead, we focused on pupil dilation, which can be modulated by sensory and noxious stimulation [13, 14], and neither differed between conditions nor correlated with connectivity effects. As also the occurrence of phosphenes or other visual sensations was not associated with connectivity effects, we conclude that our effects are unlikely to be explained by peripheral sensations. Third, as we did not obtain electrophysiological recordings during stimulation, our results cannot be generalized to online effects of tACS which might be of a different nature compared to aftereffects.

In contrast to connectivity, tACS aftereffects in power can robustly be studied at sensor level. Furthermore, optimization of high-definition tACS montages, as used here to minimize power differences between conditions, is not necessarily required to investigate aftereffects in power, although high-definition montages can in general help to spatially steer the stimulating fields and to minimize phosphenes. Stimulation outlasting power effects have been described in several recent studies [18, 19, 20, 21] and may last up to 70 min or even longer [20]. We find power effects between JP and the other stimulation conditions (IP and AP), which might be related to IP and AP stimulation being spectrally more focused on 10 Hz than JP stimulation (Supplementary Material A). Interestingly, these power effects started at about 200 s after stimulation offset. Exactly this initial initial interval of 3 min did also not lead to significant power effects in a study comparing *α*- to sham-tACS [20].

Without knowledge on immediate effects of tACS, how can stimulation outlasting changes in functional connectivity be explained? While entrainment echoes might decay within few cycles, plasticity could be crucial for longer-lasting effects. Discrimination between entrainment and plasticity was discussed recently for aftereffects in *α*-power: Vossen et al. [19] argued that characteristics of *α*-power aftereffects did not support the hypothesis of entrainment echoes, as, for example, aftereffects did not depend on phase-continuity of the tACS stimulus. Rather, mechanisms such as spike-timing-dependent plasticity (STDP) could play a role, a suggestion which was supported by Wischnewski et al. [40] via application of the N-methyl-D-aspartate receptor (NMDAR) antagonist dextromethorphan. The NMDAR antagonist, limiting excitatory STPD, abolished tACS aftereffects on *β*-power that were seen in a placebo control [40]. Similarly, Zäahle et al. [18] explained their *α*-power aftereffects by STDP.

In our study, it seems hard to delineate a mechanism explaining connectivity effects lasting for tens of seconds after tACS offset. Similar to Ahn et al. [21], we did not find a correlation between the match of stimulation frequency to the IAF and effect sizes (Supplementary Material E). We neither detected an interaction of effects with baseline power or connectivity, thus could not find indications for ongoing activity to strongly influence the efficacy of tACS. Instead, we speculate that tACS might slightly increase or decrease the probability for spikes in a certain phase of the 10 Hz cycle, potentially without strong dependence on average ongoing activity. The phase relation between the stimulating fields would then bias the probability of simultaneous spikes, directly affecting STDP of synapses. This interpretation is consistent with our finding that JP stimulation, constantly varying the relative phase of fields, showed intermediate or low connectivity changes. To directly test for the proposed mechanism, usage of NMDAR antagonists [40] might be helpful in future studies.

In conclusion, we were able to modulate phase coupling, outlasting the stimulation period, between distant brain regions by transcranial electric stimulation. Our results lend strong support to the efficacy of tACS for frequency- and space-specific modulation of neural dynamics. Based on these findings, we suggest the use of bifocal high-definition tACS to manipulate large-scale corticocortical coupling in experimental settings. In addition, clinical studies with the aim of modulating pathological or maladaptive coupling could benefit from both specificity as well as noninvasive and easy applicability of tACS.

**Table 1:** Abbreviations.

|  |  |
| --- | --- |
| tACS | transcranial alternating current stimulation |
| IP | in-phase |
| AP | anti-phase |
| JP | jittered-phase |
| EEG | electroencephalogram |
| MEG | magnetoencephalogram |
| ECG | electrocardiogram |
| EOG | electrooculogram |
| IC(A) | independent component (analysis) |
| eLORETA | exact low resolution brain electromagnetic tomography |
| AAL | automated anatomical labeling |
| SSD | spatio-spectral decomposition |
| K-S | Kolmogorov-Smirnov |
| STDP | spike-timing-dependent plasticity |
| NMDAR | N-methyl-D-aspartate receptor |
| IAF | individual alpha frequency |

## Acknowledgments

This work has been supported by DFG, SFB 936/A3 and SPP 1665/EN 533/13-1. We thank Peter Käonig, Till Schneider, Marina Fiene, Jan-Ole Radecke, and Darius Zokai for helpful discussions; Marina Fiene for proofreading of the manuscript; and Karin Deazle for technical assistance.

## Supplementary Material

### A: Properties of the stimulating fields at 5 mm resolution

We simulated the stimulating cortical electric field dependent on the stimulation condition at 5 mm spatial resolution (Fig. 1C). For JP stimulation, the phase relation of the stimulating currents constantly varied, and thus the stimulating electric field changed between the ones computed for IP and AP stimulation. As stimulating fields were maximally different between IP and AP stimulation, we restricted ourselves to reporting differences between these two conditions. Ideally, stimulation conditions should differ only in the direction or temporal phase of the applied field and not in their absolute strength or focality [38]. Our stimulation configuration was chosen to optimally meet these criteria. Given equal impedances of all four outer electrodes to the center electrode of a montage, at each voxel, the absolute field strength difference between IP and AP stimulation was below 70.0 mV/m; the average difference was −1.3 mV/m. Compared to peak field strength values of 356.1 mV/m (IP) and 369.0 mV/m (AP; 3.6 % more than IP), condition differences were thus low. In contrast, vectorial differences peaked with 637.7 mV/m and had average values of 39.6 mV/m.

Varying impedances of the tACS electrodes within one montage can affect those field distributions. In particular, our experimental procedure allowed for up to 20% difference in single impedances relative to the average impedance of the four electrodes. Maximum leakage currents between the montages are expected to occur when impedances of medial tACS electrodes are low compared to the impedances of lateral tACS electrodes. Thus, redistribution of absolute field strength was simulated for a “worst case scenario” of 50% deviation of all impedances from their average, where medial electrodes had the lowest impedances (for example: 25 *k*Ω for all medial electrodes and 75 *k*Ω for all lateral electrodes; these are much higher differences than observed in any participant). This worst case scenario led to a maximum difference in absolute field strength between IP and AP stimulation of 128.3 mV/m and an average difference of −11.3 mV/m. Although these values are higher than the field distributions assuming equal impedances, they are still small compared to peak (657.6 mV/m) and average (48.4 mV/m) vectorial differences. We therefore conclude that even in the case of unevenly distributed impedances of tACS electrodes, only a minor part of the electric field was redistributed in space between stimulation conditions.

### B: Details on stimulation conditions

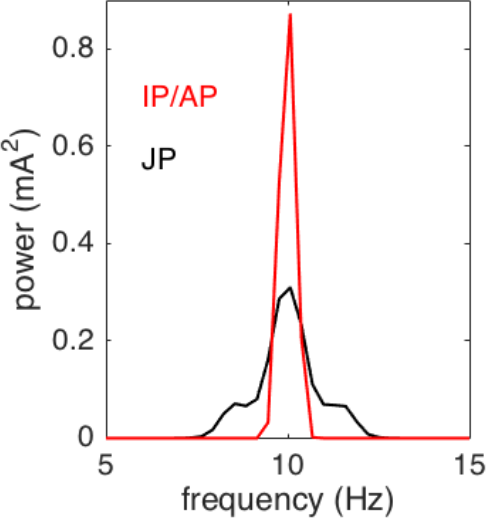

While IP as well as AP stimulation consisted of pure sinusoidal currents, frequency “sweeps” were independently included in both stimulating currents for the JP condition to change the phase relationship of the stimulating fields over time. The sweeps included several linear shifts in frequency between 9.5 and 10.5 Hz. Consequently, power of the stimulating current peaked at 10 Hz, but also extended to the neighboring frequencies. In contrast, sinusoidal stimulation used for IP and AP stimulation showed a sharp peak in power at 10 Hz. Coherence between JP stimulation currents was below 0.02 for all frequencies.

### C: Power and connectivity time courses

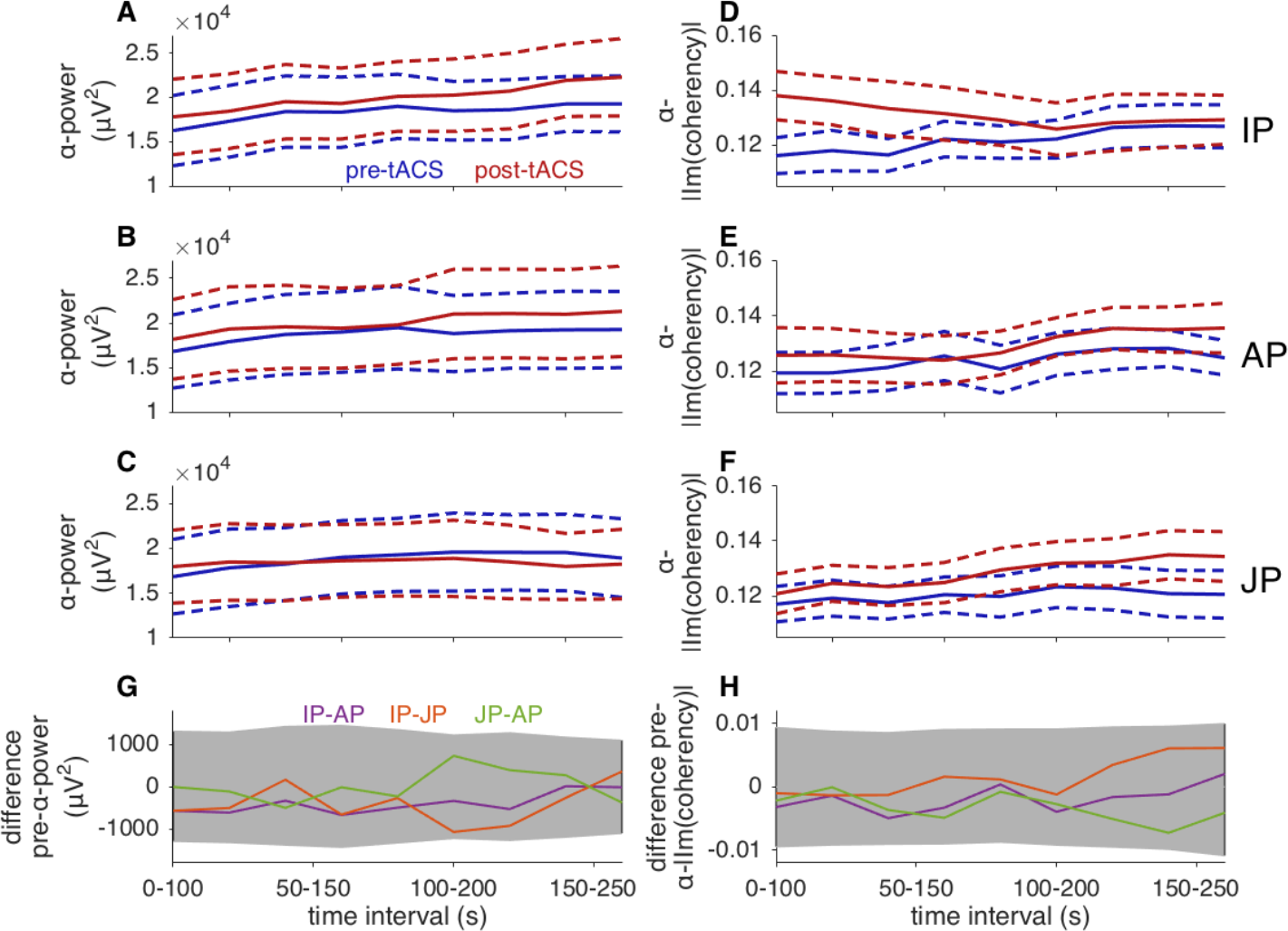

Grand average power (A-C) and imaginary coherence in the *α*-band (D-F) shown separately for IP (A, D), AP (B, E) and JP (C, F) stimulation as well as for pre-(*blue*) and post-tACS (*red*) intervals. Dashed lines indicate the mean ± standard error of the mean. Direct pre-post comparisons of connectivity in single conditions are confounded by order effects and by *α*-power increasing over time. Baseline condition differences in power (G) and connectivity (H) were not significant (uncorrected *p* > 0.05 for all time windows, permutation test).

### D: Initial decay of connectivity effects in 2 s time steps

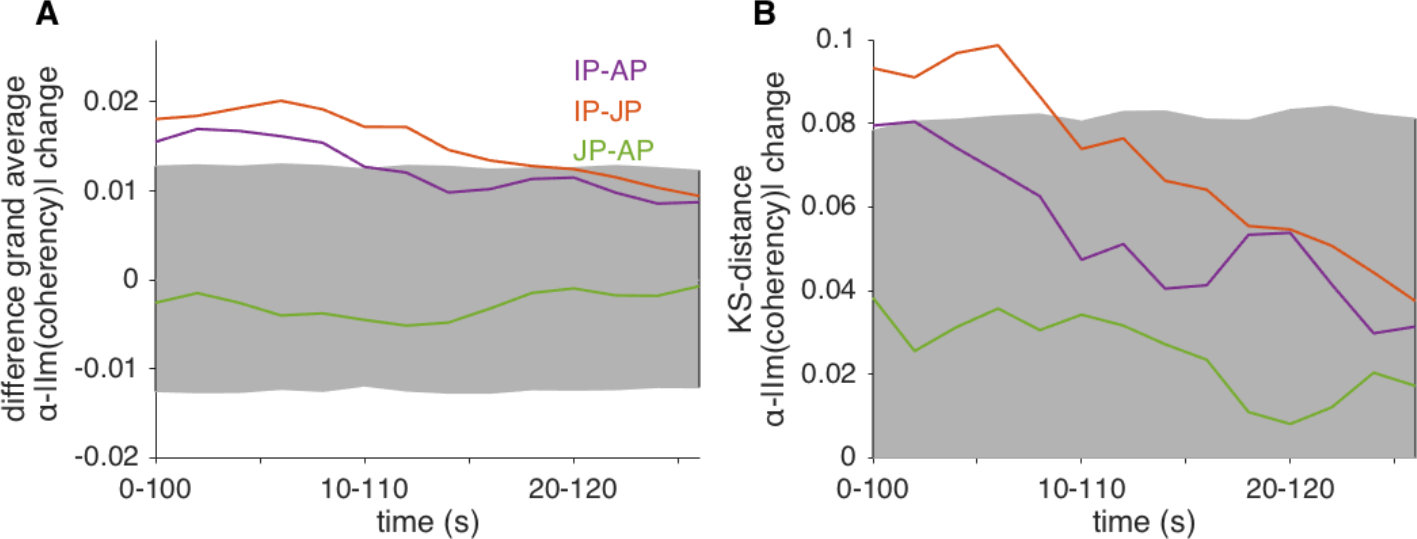

Differences in grand average change in imaginary coherence (A) and corresponding KS-distances (B). While both measures were computed within windows of 100 s as in Fig. 2, segments were shifted in 2 s steps to describe the initial decay of effects. Effects gradually decayed with the time lag. All effects lost significance before time window 20-120 s.

### E: Connectivity effects correlate neither with power effects nor with parameters of individual physiology

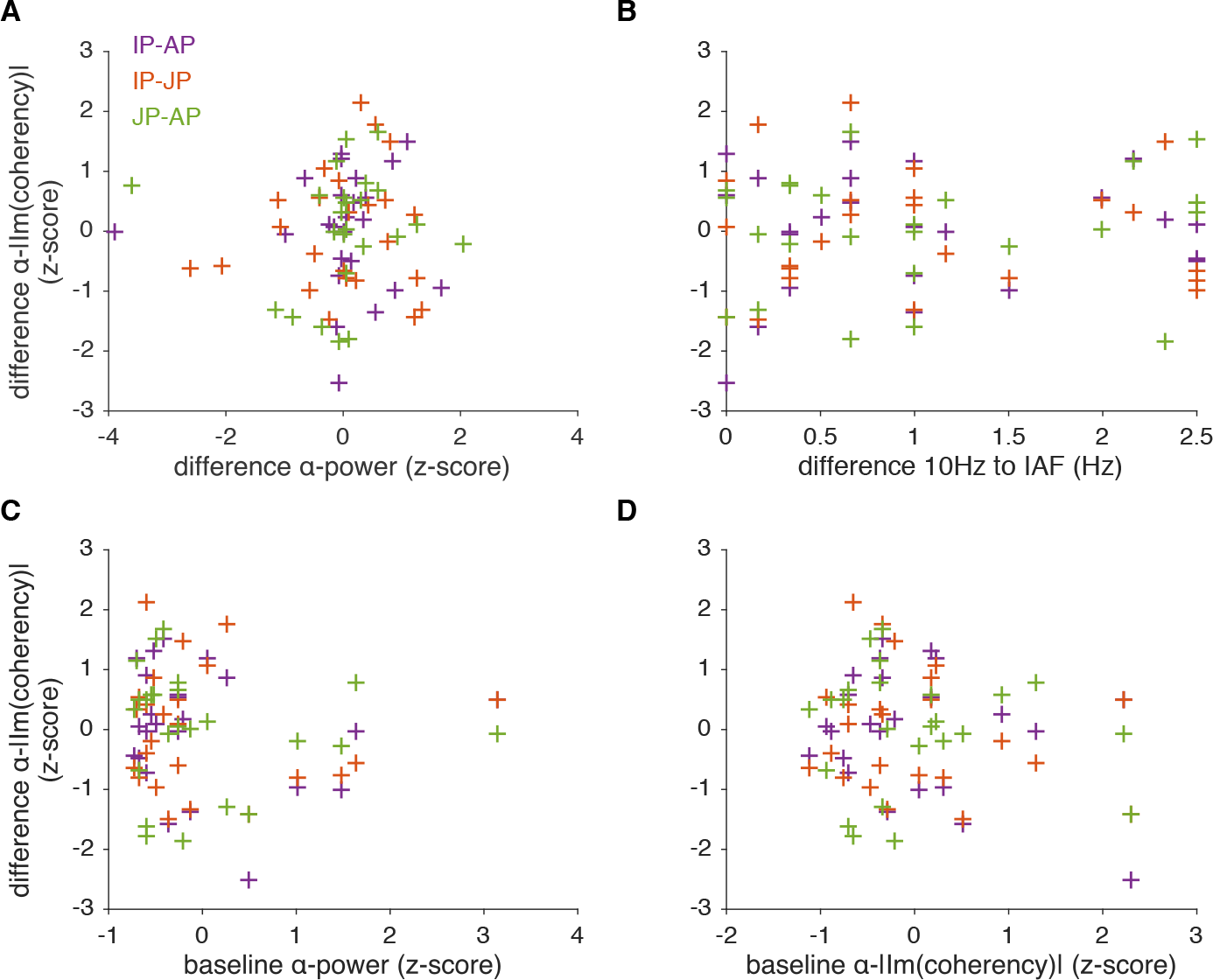

Pearson correlations coefficients over participants between the main effect — the change in grand average *α*-imaginary coherence between tACS conditions within the time interval 0-100 s — and other metrics, namely: A: change in grand average *α*-power between tACS conditions, within the same time interval; B: difference between stimulation frequency (10 Hz) and individual alpha frequency (IAF); C: baseline *α*-power of all pre-recordings; D: baseline *α*-imaginary coherence of all pre-recordings. None of the correlations reached significance (uncorrected *p* >0.05 for all condition contrasts). Imaginary coherence and power values were z-scored. *α* denotes the stimulated frequency band (9-11 Hz).

### F: Connectivity matrix of average α imaginary coherence change, contrasting IP and AP stimulation conditions, without thresholding at *p*<0.05

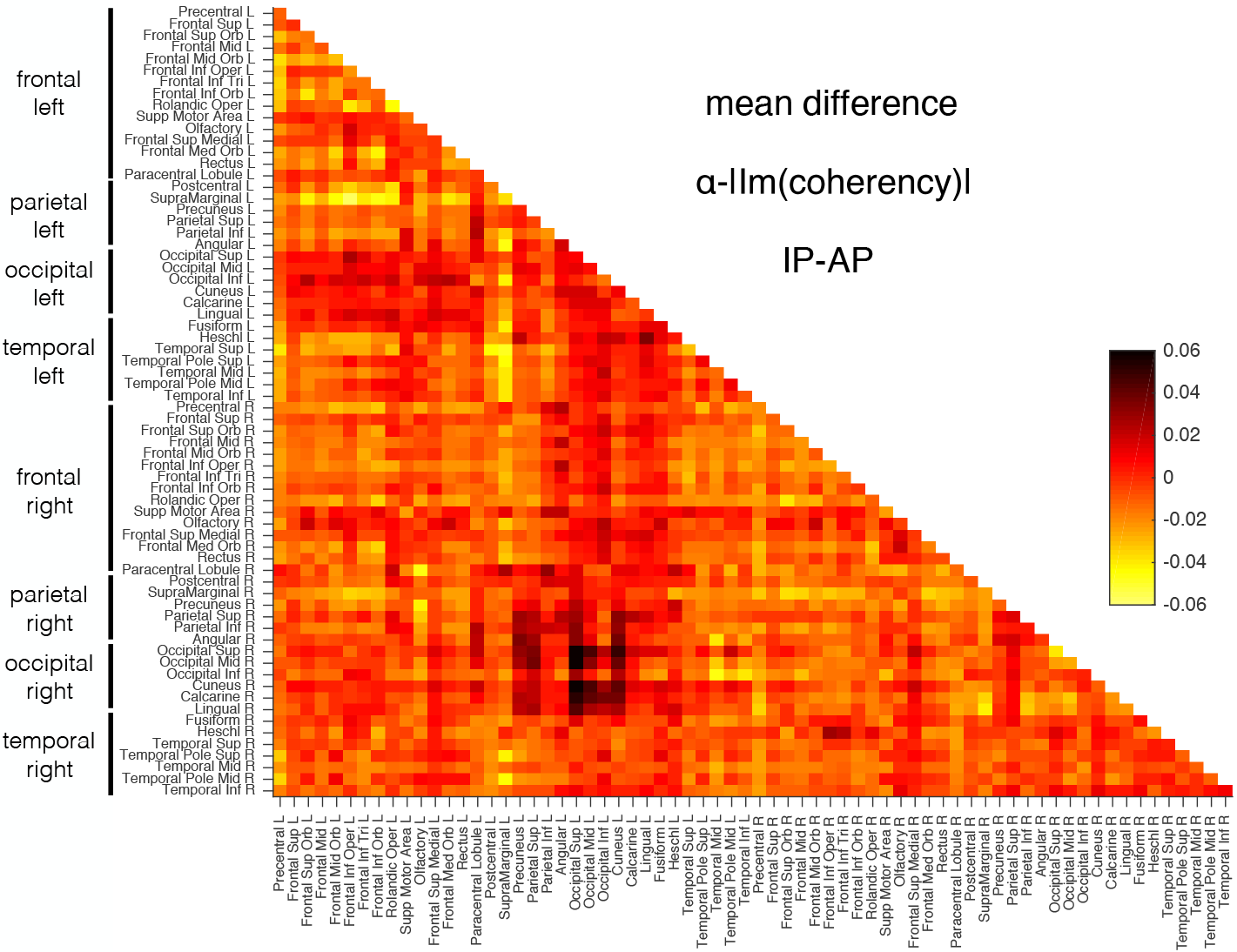

Interhemispheric coupling change between the stimulated occipito-parietal regions shows largest condition differences.

### G: Pupil dilation, occurrence of visual sensations and relation to connectivity effects

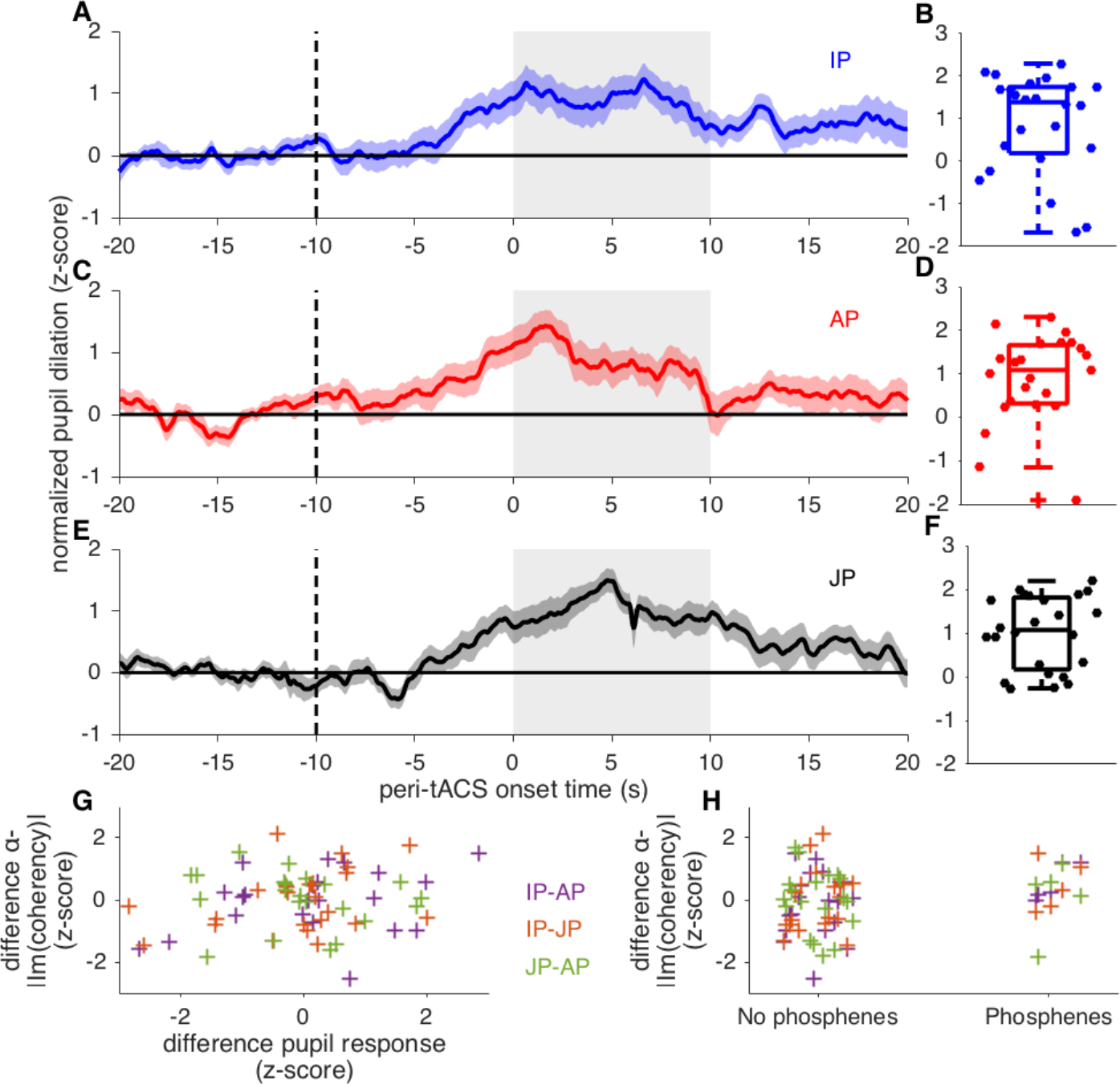

Grand average z-scored and normalized pupil dilation for IP (A), AP (C) and JP (E) stimulation relative to tACS onset. While ramp-in of tACS started at time −10 s (dashed vertical line), the current reached its maximum amplitude at time zero. In all three conditions, there was a clear increase in pupil dilation around tACS onset which was largest within the first 10 s (gray shaded area). Mean values of pupil dilation within this interval (“responses”, distributions across participants are shown in panels B, D, F) did not differ significantly between conditions (uncorrected *p* >0.5, Wilcoxon signed rank sum test). Condition differences between responses did furthermore not correlate with effects (G; uncorrected *p* > 0.1 for all condition contrasts). The five participants with visual perceptions such as phosphenes did not show significantly stronger effects than the remaining participants without visual perceptions (H; uncorrected *p* > 0.1 for all condition contrasts, Wilcoxon rank sum test).

